# Susceptibility and cellular responses of COPD airway epithelium to cigarette smoke in stem cell-derived ALI models

**DOI:** 10.64898/2026.10.27.762365

**Authors:** Jialei Jiang, Yushan Tian, Yinuo Zhao, Chen Tan, Xiao Li, Hongjuan Wang, Longjiang Fan, Hongyu Chen

**Author notes:** Corresponding authors: Hongyu Chen and Longjiang Fan. These authors contributed equally (Jialei Jiang, Yushan Tian, Yinuo Zhao).

## Abstract

Despite cigarette smoke being the chief driver of chronic obstructive pulmonary disease (COPD), numerous patients maintain their smoking habits by opting for low-yield or slim cigarettes under the misconception of diminished health risks. Differential effects of high-tar conventional versus low-tar slim cigarettes on COPD airway epithelium remain poorly defined. We established an air–liquid interface (ALI) model using patient-derived COPD airway epithelial stem cells and exposed cultures to high-tar conventional smoke (TC), low-tar slim smoke (SC), or filtered air (NC). Both regimens induced IL-1β inflammation, ciliary structural changes, and oxidative stress, with TC eliciting stronger responses. Single-cell RNA sequencing resolved six epithelial populations. In ciliated cells, hdWGCNA showed NC modules associated with cilium organization and assembly, SC adaptive defense responses, and TC more extensive reprogramming of ciliary functional networks. Basal cells displayed stress-pathway activation, squamous metaplasia signatures, and developmental regression toward a stem-like state, with TC producing approximately 1.7-fold stronger regression than SC. Secretory cells exhibited identity erosion, antimicrobial program downregulation, and a shift toward keratinization and squamous-oriented programs, again more pronounced under TC. These findings indicate that high-tar conventional smoke exerts greater influence than low-tar slim smoke on COPD airway epithelium across inflammatory, structural, and lineage-specific molecular dimensions, providing high-resolution insight into cell-type-specific responses to continued smoke exposure.

## Introduction

COPD persists as a widespread public health crisis, distinguished by ongoing respiratory symptoms and unrelenting, irreversible airflow reduction[1]. Cigarette smoke represents the primary environmental etiology driving both the onset and exacerbation of COPD [2]. Upon inhalation, the complex mixture of toxic and reactive chemicals in cigarette smoke directly damages the airway epithelium, inducing oxidative stress [3], cell mortality, and chronic inflammation [4], which ultimately lead to goblet cell metaplasia, mucus hypersecretion [5], and ciliary dysfunction [6].

The airway epithelium acts as the frontline physicochemical barrier against inhaled noxious agents. Under homeostatic conditions, airway basal cells function as resident stem and progenitor cells, undergoing self-renewal and controlled differentiation into ciliated and secretory cells to maintain mucociliary clearance [7]. In COPD, however, this regenerative architecture is markedly compromised due to stem cell exhaustion and aberrant lineage reprogramming [8, 9], conferring heightened susceptibility to subsequent environmental insults. Although conventional immortalized cell lines and animal models have provided foundational insights, they fail to faithfully recapitulate the cellular complexity, differentiation capacity, and disease-specific phenotypes of the human airway [10]. Consequently, patient-derived airway epithelial stem cells cultured at the air-liquid interface (ALI) have emerged as a highly translationally relevant platform to model disease-specific epithelial dynamics [11].

Despite clear clinical guidelines advocating complete smoking cessation, a substantial cohort of COPD patients continues to smoke post-diagnosis. Driven by commercial marketing and the common perception that low-tar or slim products may be less harmful, many patients switch to these alternative cigarettes rather than quitting. Nevertheless, the extent to which distinct cigarette formulations, specifically conventional high-tar versus low-tar slim cigarettes, exert differential deleterious effects on an already compromised COPD airway epithelium remains poorly defined [12]. Unraveling these persistent toxicological impacts under conditions simulating ongoing tobacco exposure is imperative for clarifying disease pathogenesis and informing public health strategies.

Furthermore, conventional bulk analytical approaches reflect population-averaged signals, thereby masking the marked cellular heterogeneity and cell-type-specific stress response pathways that drive tissue injury. Recent advances in single-cell RNA sequencing offer unprecedented resolution to dissect discrete epithelial subpopulations, stress-induced transitional states, and altered lineage trajectories in response to environmental perturbations [13–15].

In this study, we established an ALI smoke-exposure model using stem cells derived from COPD patients to systematically evaluate the susceptibility and cellular responses of diseased airway epithelium to cigarette smoke. By combining physiological assessments of oxidative damage, IL-1β-driven inflammatory signaling, MUC5AC-mediated mucin expression, and ciliary structural integrity with single-cell transcriptomic profiling across different smoke exposure conditions, including high-tar conventional and low-tar slim cigarettes, we pinpointed the cell-type-specific vulnerabilities and molecular programs that drive ongoing epithelial disruption in COPD.

## Results

### Cigarette smoke exposure drives acute inflammatory responses, ciliary impairment, and oxidative stress in COPD airway epithelium

To investigate the cytotoxic and pathological effects of cigarette smoke on already compromised airway epithelium, we established an in vitro air-liquid interface (ALI) exposure system using primary stem cells isolated from patients with COPD. Cells were expanded, differentiated on Transwell inserts under ALI conditions, and subsequently exposed to high-tar conventional cigarette smoke (TC), low-tar slim cigarette smoke (SC), or filtered air (NC) using an RM20S smoking machine integrated with continuous basolateral medium perfusion (Fig. 1A).

**Figure 1.**
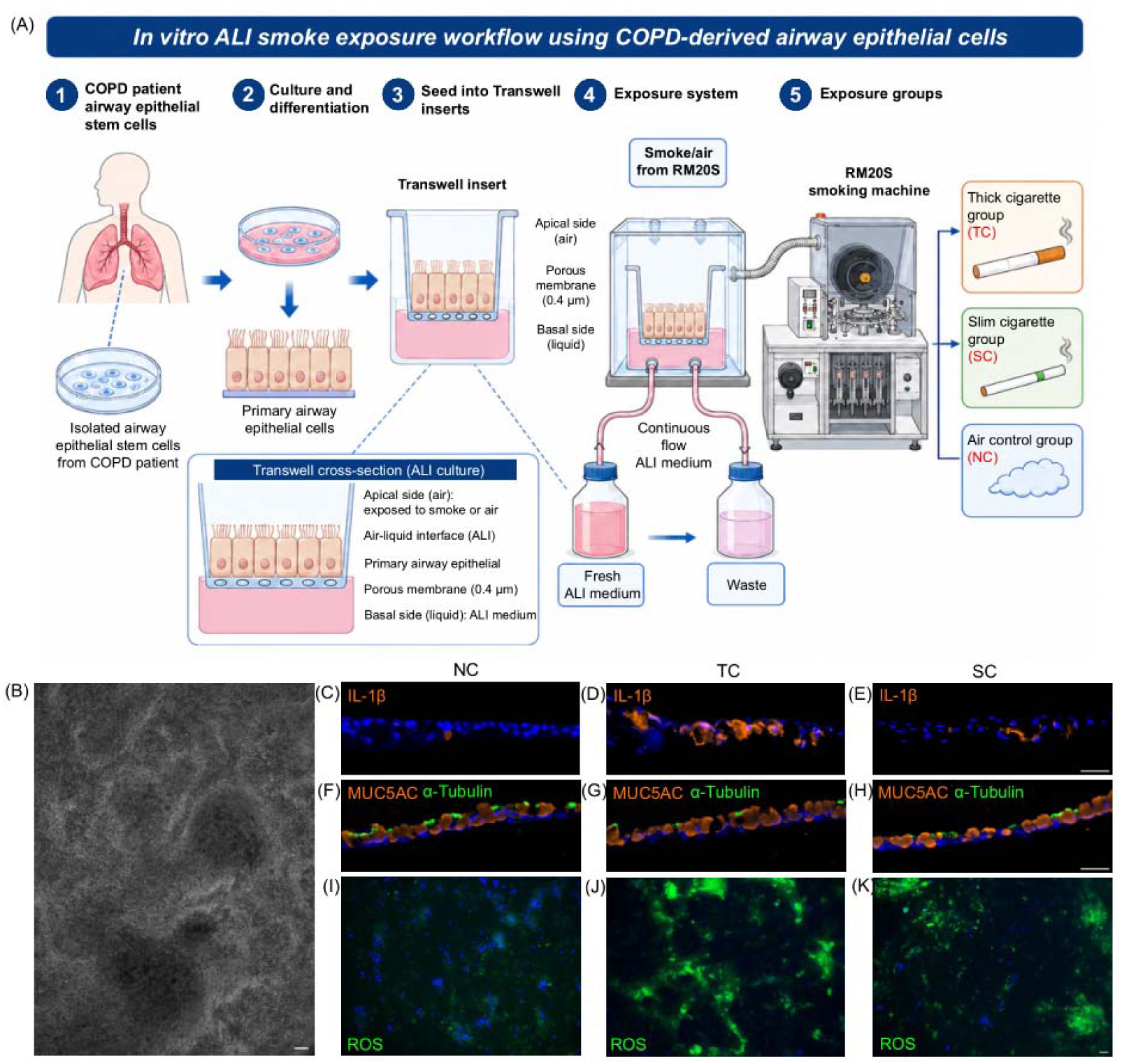
In vitro air–liquid interface smoke exposure workflow and effects of different cigarette treatments on inflammatory response, mucus-associated marker expression, and oxidative stress in COPD patient-derived primary airway epithelial cells. (A) Schematic of the in vitro air–liquid interface (ALI) smoke exposure workflow using COPD patient-derived airway epithelial stem cells. Cells were expanded, differentiated in Transwell inserts under ALI conditions, and exposed to cigarette smoke or air via an RM20S smoking machine with continuous basolateral medium perfusion.(B) Representative bright-field image of COPD patient-derived primary airway epithelial cells.(C–E) Representative fluorescence images of IL-1β in cells exposed to air control (NC), high-tar conventional cigarette (TC), or low-tar slim cigarette (SC) treatment.(F–H) Representative fluorescence images of MUC5AC and α-Tubulin following the indicated treatments. MUC5AC is shown in orange and α-Tubulin in green.(I–K) Representative merged fluorescence images of reactive oxygen species (ROS) following the indicated treatments. ROS is shown in green, and nuclei are counterstained with DAPI (blue).

Following differentiation, COPD-derived epithelial cells formed a confluent monolayer exhibiting typical pseudostratified epithelial morphology (Fig. 1B). We first evaluated smoke-induced inflammatory activation by immunofluorescence staining for IL-1β. Compared with the air control, both TC and SC exposure markedly increased IL-1β expression in the epithelium, confirming that cigarette smoke directly triggers acute airway inflammation. Quantification of fluorescence intensity showed a pronounced elevation in IL-1β signal following TC exposure, whereas SC exposure resulted in a relatively moderate increase (Fig. 1C-E).

We next examined the impact of smoke exposure on epithelial cell lineage markers through dual immunofluorescence staining for MUC5AC and α-Tubulin. In air-treated controls, prominent α-Tubulin-positive ciliary structures were readily detected across the apical surface. Cigarette smoke exposure compromised ciliary integrity, with TC causing a severe reduction in α-Tubulin signal, while SC-treated tissues preserved a higher fraction of ciliated structures (Fig. 1F-H). Interestingly, strong MUC5AC immunoreactivity was persistently observed across all groups with only modest inter-group variations, underscoring the intrinsic hypersecretory baseline characteristic of COPD-derived epithelium.

Finally, epithelial oxidative stress was assessed via ROS fluorescence. Whereas air control tissues displayed minimal basal ROS, both cigarette smoke regimens elicited a clear elevation in intracellular ROS generation, reflecting pervasive smoke-induced oxidative damage. Specifically, TC exposure induced a widespread and intense ROS signal, while SC exposure produced a more moderate increase in ROS levels (Fig. 1I-K).

### Single-cell transcriptomic profiling reveals cellular heterogeneity and distinct epithelial subpopulations in COPD ALI cultures

To resolve the cellular heterogeneity and dissect the transcriptomic dynamics of COPD-derived airway epithelium following cigarette smoke exposure, we performed single-cell RNA sequencing (scRNA-seq) on cells harvested from the air-liquid interface (ALI) cultures across the air control (NC), low-tar slim cigarette (SC), and high-tar conventional cigarette (TC) groups (Fig. 2A). Quality control metrics-including the distribution of detected genes (nFeature_RNA), unique molecular identifiers (nCount_RNA), and mitochondrial gene percentages-exhibited consistent profiles across all individual biological replicates and annotated cell clusters (Fig. S1A, B), confirming the robustness and high quality of the single-cell library preparation. Unsupervised graph-based clustering and UMAP visualization resolved the major epithelial lineages and transitional states within the cultured tissues (Fig. 2B, C). Based on canonical cell-type-specific markers, the captured cells were categorized into six major populations: Basal cells (KRT5, TP63), Basal-suprabasal provisional cells (KRT17, SERPINB3), Secretory cells (SCGB1A1, SCGB3A1, MUC5AC), Secretory-like provisional cells (LY6D, SPINK5), Ciliated cells (FOXJ1, TPPP3, DNAH5), and a distinct cluster of Stressed cells marked by the prominent upregulation of stress-response genes (HSPA1A, DDIT3, JUN) (Fig. 2B-E).

**Figure 2.**
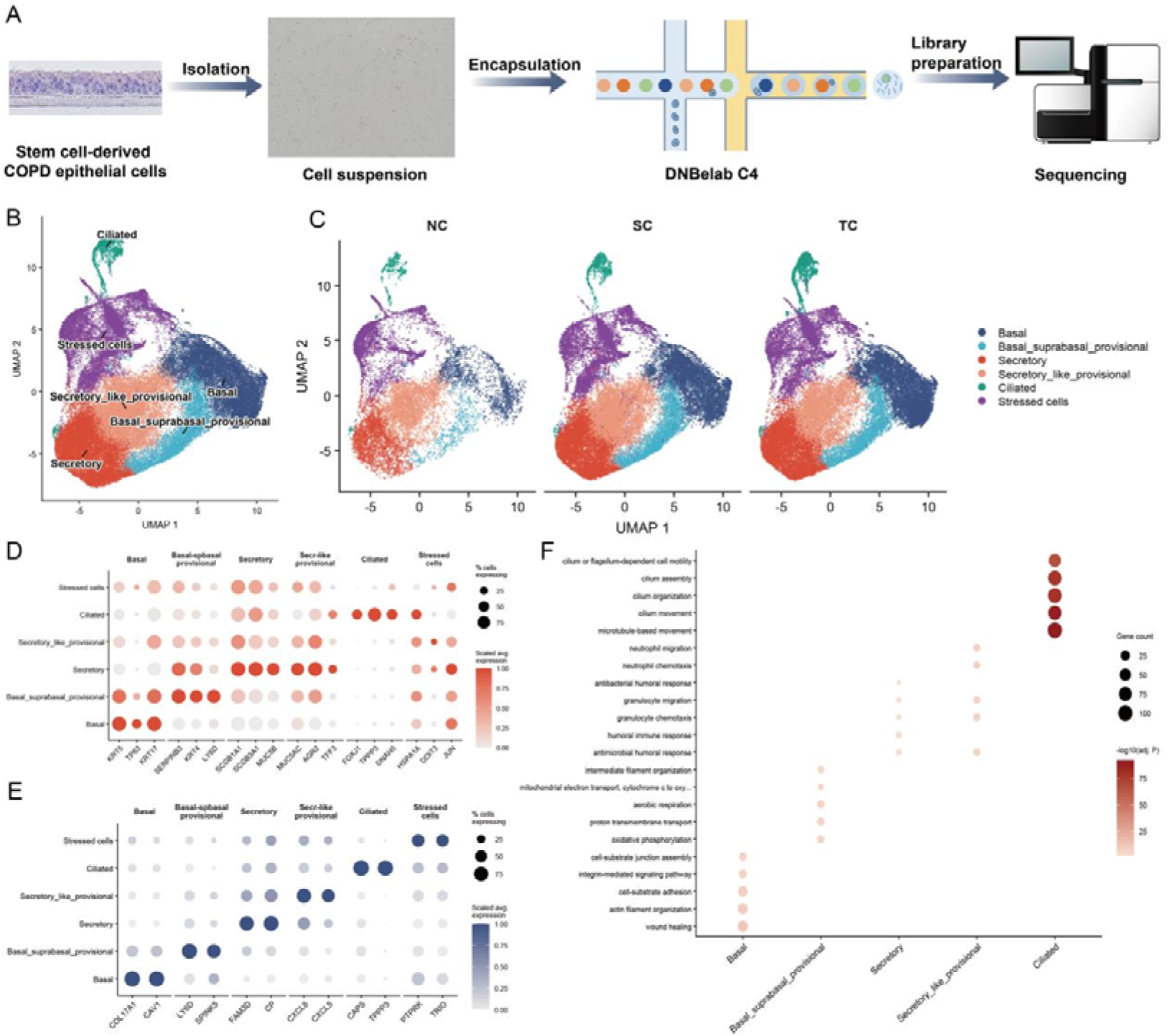
Single-cell transcriptomic profiling of COPD patient-derived airway epithelial cells cultured at the air–liquid interface (ALI) under different cigarette smoke exposure conditions. (A) Overview of the single-cell RNA sequencing experimental workflow, spanning cell isolation, suspension preparation, encapsulation via the DNBelab C4 platform, library preparation, and high-throughput sequencing. (B) UMAP projection of all sequenced cells, colored and annotated by major cell types and provisional states, including Basal, Basal_suprabasal_provisional, Secretory, Secretory_like_provisional, Ciliated, and Stressed cells. (C) UMAP plots displaying the distribution of cells across the air control (NC), low-tar slim cigarette (SC), and high-tar conventional cigarette (TC) exposure groups. (D) Dot plot showing the percentage of expressing cells (dot size) and scaled average expression (color intensity) of canonical lineage and marker genes across identified cell types. (E) Dot plot illustrating the expression patterns of additional functional and subpopulation-specific marker genes across distinct epithelial clusters. (F) Gene Ontology (GO) biological process enrichment analysis of marker genes across major epithelial cell populations, with dot size representing gene count and color intensity indicating statistical significance.

The expression profiles of canonical lineage markers confirmed the identity of each subpopulation (Fig. 2D). Specifically, basal and provisional populations robustly expressed structural and stemness markers (e.g., KRT5, TP63, KRT17), secretory groups enriched classic mucin and club cell markers (e.g., SCGB1A1, MUC5AC), and ciliated clusters showed high expression of motility-related genes (e.g., FOXJ1, TPPP3, DNAH5) (Fig. 2D). Furthermore, cell-type-specific functional markers-including COL17A1 and CAV1 in basal compartments, and CXCL8, CXCL5, CAPS, and PTPRK across differentiated layer-further delineated the transcriptional specialization among subpopulations (Fig. 2E). Gene Ontology (GO) enrichment analysis of cell-type-specific markers revealed distinct functional programs aligned with individual epithelial identities (Fig. 2F). Ciliated cells were heavily enriched in biological processes governing “cilium or flagellum-dependent cell motility” and “cilium assembly,” whereas secretory and provisional populations showed distinct functional signatures related to defense responses and structural organization (Fig. 2F). These single-cell transcriptomic data establish a high-resolution cellular baseline of the COPD airway epithelial model, providing a foundation for investigating smoke-induced cellular reprogramming and subtype-specific vulnerabilities.

### hdWGCNA reveals distinct modular functional reprogramming of ciliated cells under TC and SC exposure

To dissect the cell-type-specific transcriptomic responses of ciliated cells to different cigarette smoke regimens, we performed hdWGCNA on the Ciliated cell population. This approach enabled the unbiased identification of co-expression modules that capture coordinated gene programs associated with air control (NC), low-tar slim cigarette (SC), or high-tar conventional cigarette (TC) exposure.

The hdWGCNA dendrogram resolved multiple robust co-expression modules (Fig. 3A). Module feature gene expression patterns across treatment groups revealed clear exposure-specific distributions. Specifically, the turquoise and brown modules were predominantly expressed in the NC group, whereas the green module was specifically enriched under SC exposure. The yellow, blue, pink, and magenta modules were markedly elevated in the TC group. By contrast, the red module was shared across both SC and TC conditions, exhibiting higher overall expression under TC exposure (Fig. 3B). Hub gene interaction networks further delineated these co-expression modules into discrete functional groupings (Fig. 3C). The brown and turquoise modules formed the NC-related network, green constituted the SC-related network, red represented a shared SC and TC smoke response, and yellow, blue, pink, and magenta together defined the TC-specific functional network.

**Figure 3.**
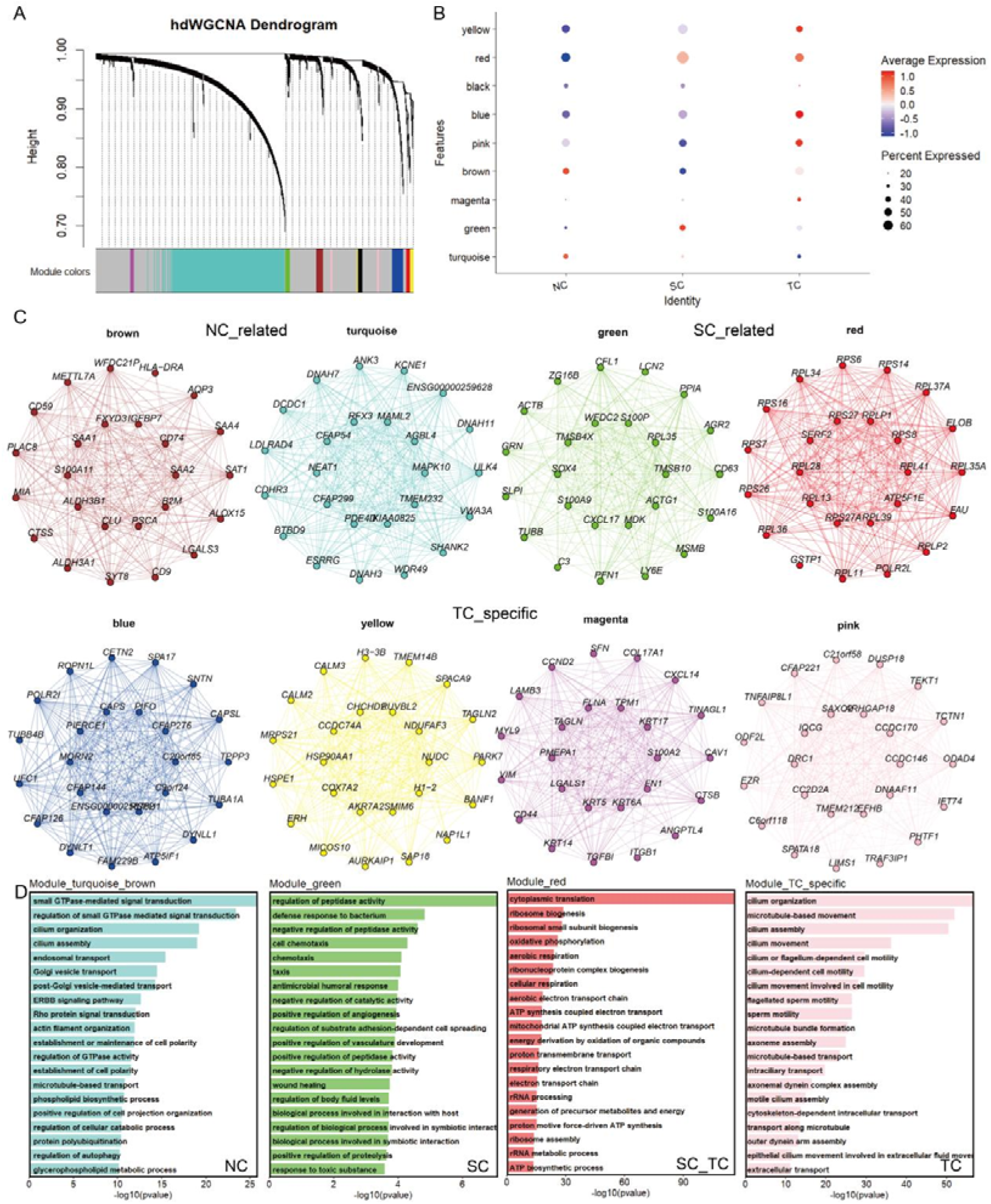
hdWGCNA analysis of ciliated cells reveals exposure-specific co-expression modules and functional programs in COPD airway epithelium. (A) Dendrogram of the hdWGCNA co-expression network constructed from Ciliated cells, with modules indicated by color bars below. (B) Dot plot showing average expression and percent expressed of module feature genes across different groups. (C) Hub gene networks of the identified modules. NC-related modules (brown and turquoise), SC-related module (green), shared SC/TC module (red), and TC-specific modules (yellow, blue, pink and magenta) are shown. (D) Gene Ontology biological process enrichment of genes from NC-related modules, SC-related module, shared SC/TC module, and TC-specific modules.

Gene Ontology (GO) biological process enrichment analysis of the module genes clarified their underlying functional pathways (Fig. 3D). NC-related modules (turquoise and brown) were strongly enriched for processes governing small GTPase-mediated signal transduction, cilium organization, cilium assembly, and cell polarity establishment, reflecting homeostatic ciliary maintenance in unexposed cells. The SC-related module (green) enriched pathways involved in defense responses to bacteria, chemotaxis, antimicrobial humoral responses, and wound healing, consistent with moderate adaptive and host defense programs. The shared red module was predominantly enriched for cytoplasmic translation, ribosome biogenesis, oxidative phosphorylation, and aerobic respiration, pointing to widespread translation and metabolic demands under smoke stress. In contrast, TC-specific modules were heavily enriched for cilium organization, microtubule-based movement, cilium assembly, and axoneme assembly (Fig. 3D), indicating a profound, loss-of-function or compensatory remodeling of the core ciliary apparatus under high-tar conventional smoke exposure.

### Cigarette smoke exposure drives extensive transcriptional reprogramming and developmental regression in basal lineage cells

To systematically evaluate the transcriptomic perturbations induced by cigarette smoke within the basal epithelial lineage, we analyzed condition-level differentially expressed genes (DEGs) across the air control (NC), low-tar slim cigarette (SC), and high-tar conventional cigarette (TC) groups. Differential expression analysis revealed that both TC and SC exposures triggered thousands of DEGs in both Basal and Basal_suprabasal_provisional cell populations, displaying a highly concordant direction of change between the two smoke regimens (Fig. 4A).

**Figure 4.**
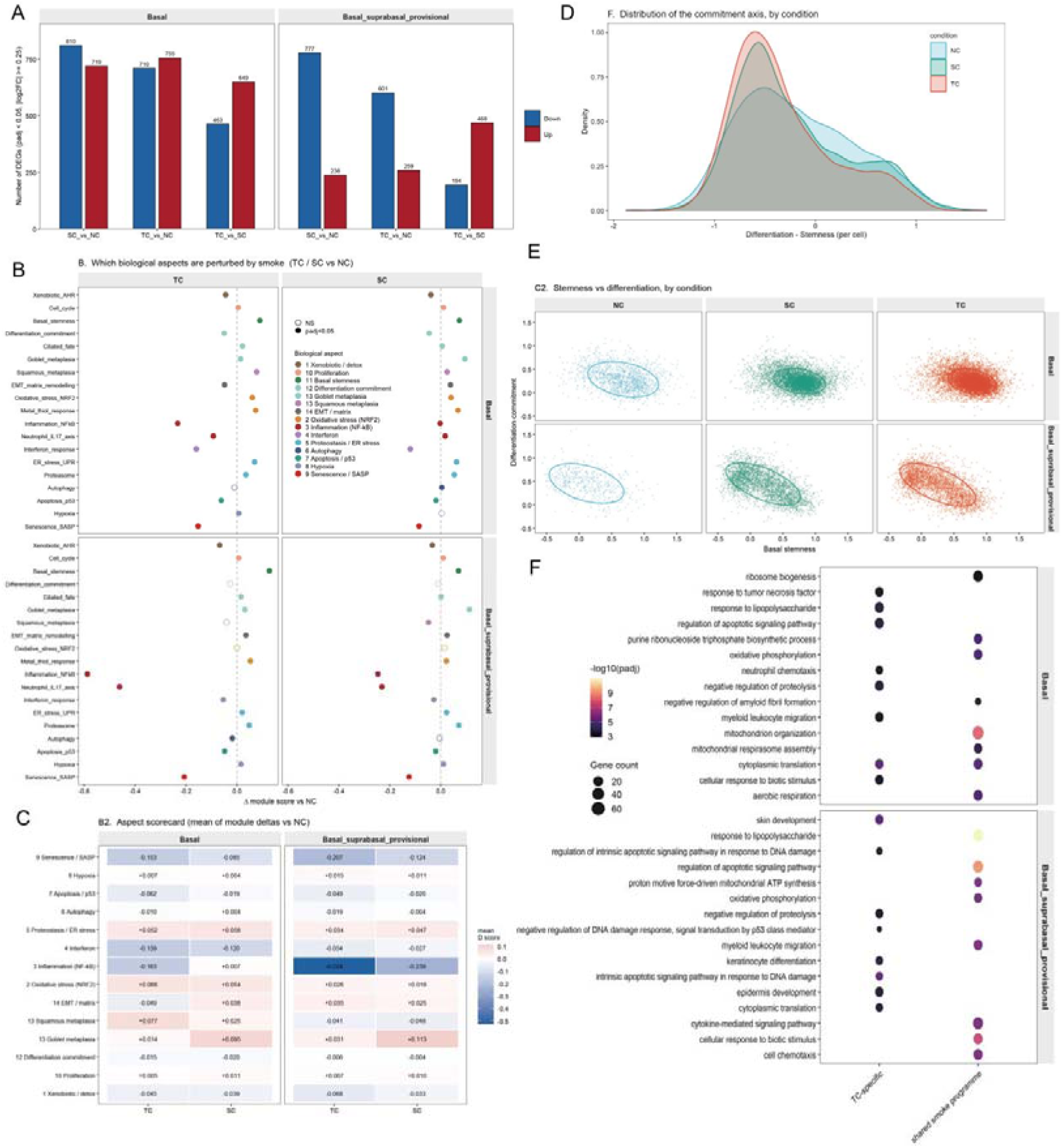
Condition-level transcriptomic perturbations, pathway scorecarding, and developmental regression of basal lineage cells in response to cigarette smoke exposure. (A) Bar plots showing the number of degs across treatment comparisonsin Basal and Basal_suprabasal_provisional cell populations. (B) Dot plots illustrating the perturbation of 18 COPD- and smoke-related biological modules across treatments, where dot color indicates upregulation (red) or downregulation (blue), and filled/open circles denote statistical significance. (C) Aspect scorecard summarizing the mean module score changes across biological themes for Basal and Basal_suprabasal_provisional cells under TC and SC exposures. (D) Density distribution plot showing the shift along the commitment (differentiation vs. stemness) axis across NC, SC, and TC conditions. (E) Scatter plots correlating stemness versus differentiation scores across conditions, demonstrating smoke-induced transcriptional shifting toward a stem-like state. (F) GO enrichment analysis of the shared smoke response program.

Aspect scorecard and modular enrichment analyses demonstrated that cigarette smoke robustly upregulated stress-associated and pathological remodeling pathways, including NRF2-mediated oxidative stress, metal-thiol responses, endoplasmic reticulum stress/unfolded protein response (ER stress/UPR), and squamous metaplasia (particularly prominent in Basal cells), while suppressing interferon and senescence-associated secretory phenotype (SASP) programs (Fig. 4B, C). Notably, classical inflammatory pathways exhibited a downward trend in single-cell mRNA levels across smoke-exposed groups, a phenomenon that likely reflects survivor bias following acute toxic injury or active transcriptional negative feedback regulation (Fig. 4B, C).

Crucially, developmental trajectory and composite index analyses uncovered a profound disruption of normal epithelial differentiation dynamics. Rather than accelerating forward maturation, cigarette smoke shifted cells backward along the normal differentiation axis, pulling pre-committed suprabasal cells back toward a stem-like, less-committed state (Fig. 4D, E). Projection onto the NC-derived differentiation axis confirmed this de-commitment and differentiation arrest, with TC exerting a more pronounced regression effect-approximately 1.7-fold that of SC (Fig. 4H, I). This transcriptomic regression precisely mirrors the classic pathological hallmarks of COPD, namely basal cell hyperplasia and differentiation arrest. Furthermore, quantitative comparison revealed that low-tar slim cigarettes (SC) replicated the vast majority of the transcriptional response direction of conventional cigarettes (TC), mitigating developmental regression by only about 40%. This finding directly challenges commercial marketing claims regarding the safety profile of alternative low-tar products, providing critical evidence that reduced-tar formulations fail to eliminate underlying developmental and cellular toxicity.

### Secretory lineage cells undergo smoke-driven identity erosion and squamous-oriented reprogramming, with stronger effects under TC exposure

To determine how cigarette smoke perturbs the secretory epithelial compartment in the COPD ALI model, we analyzed secretory and secretory-like provisional cells across NC, SC, and TC conditions (33,186 cells total). Both smoke regimens induced extensive transcriptional changes dominated by downregulation versus air control. Secretory cells showed 4,030 DEGs (SC vs NC: 169 up / 3,861 down) and 4,406 DEGs (TC vs NC: 233 up / 4,173 down); the direct TC versus SC comparison yielded 819 DEGs. Secretory-like provisional cells exhibited smaller but substantial responses (SC vs NC: 1,319; TC vs NC: 1,366; TC vs SC: 823 DEGs) (Fig. 5A). The predominance of downregulated genes partly reflected the elevated inflammatory baseline already present in unexposed COPD epithelium. Gene Ontology analysis revealed a consistent shift toward keratinization and squamous programs. Shared upregulated pathways included keratin filament, cornified envelope, intermediate filament organization, and cytokine/chemokine activity. TC-specific pathways further emphasized collagen-containing extracellular matrix, response to lipopolysaccharide, non-canonical NF-κB signaling, and apoptotic processes, whereas SC-specific sets yielded few significant enrichments (Fig. 5B).

**Figure 5.**
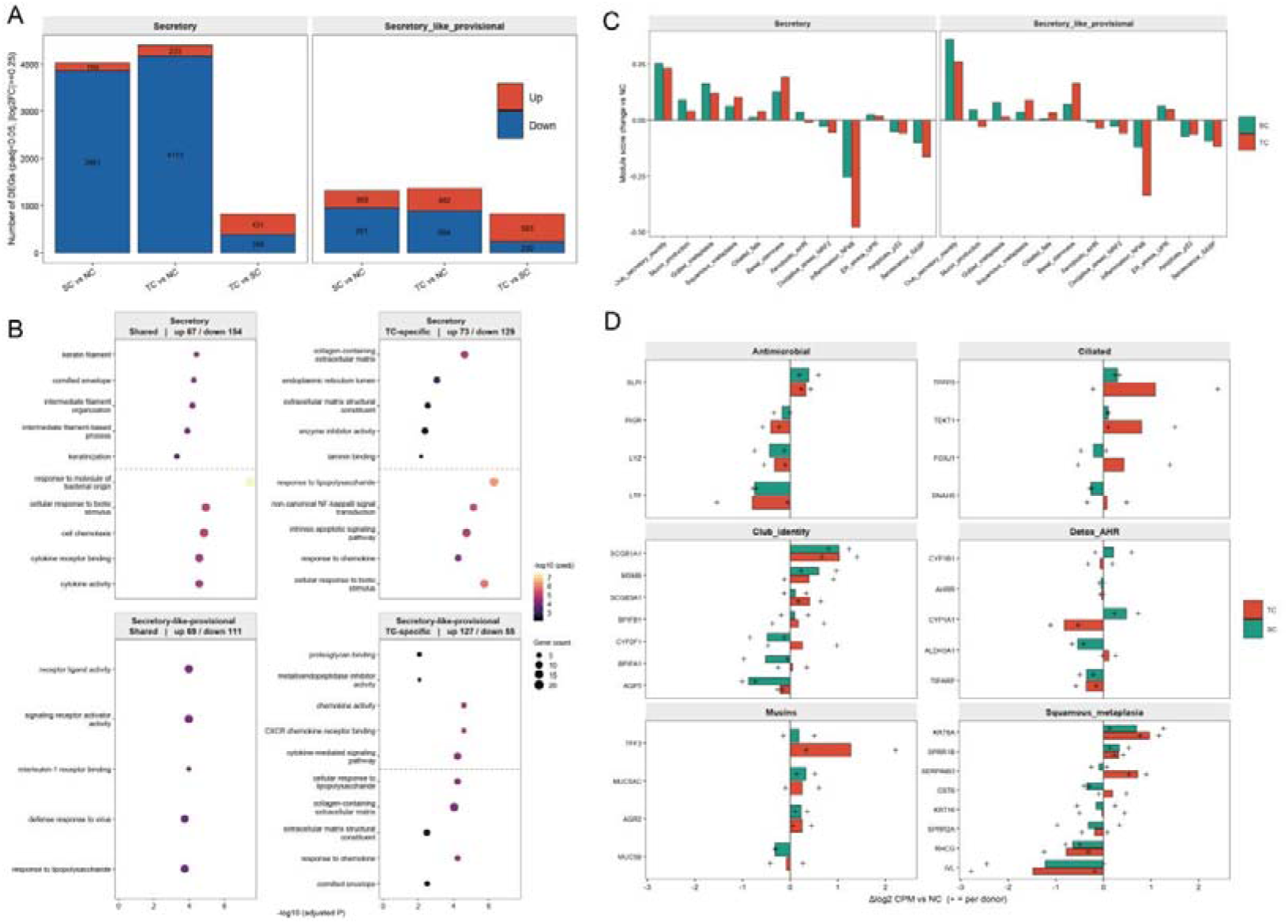
Transcriptional reconstruction and developmental axis perturbation of the Secretory lineage under different exposure conditions. (A) Number of differentially expressed genes (DEGs) (up-regulated in red, down-regulated in blue) in secretory and secretory-like provisional cells across different pairwise comparisons. (B) GO biological process enrichment analysis of shared and condition-specific DEGs in secretory and secretory-like provisional cells following smoke exposure (upper block of each panel: up-regulated genes; lower block: down-regulated genes). (C) Changes in core functional module scores relative to the NC baseline across SC (green) and TC (red) conditions. (D) Pseudobulk quantitative expression changes of representative marker genes across major functional panels, including antimicrobial proteins, ciliated markers, club cell identity, detoxification/AHR pathways, mucins, and squamous metaplasia markers.

Module-score analysis confirmed elevation of Club_secretory_identity, Goblet_metaplasia, and Squamous_metaplasia scores under both conditions, with Squamous_metaplasia more pronounced under TC; Inflammation_NFκB and Senescence/SASP scores declined (Fig. 5C). Gene-level quantification showed broad downregulation of antimicrobial genes (SLPI, PIGR, LYZ, LTF), mixed Club-identity changes, TC-biased induction of TFF3, and stronger upregulation of squamous markers (KRT6A, SPRR1B, SERPINB3, CST6, KRT16) under TC than SC. Terminal cornification markers declined, consistent with early rather than completed squamous differentiation (Fig. 5D). Projection onto an NC-defined Secretory-to-provisional axis demonstrated compression of the continuum: mature Secretory cells shifted toward the provisional pole while provisional cells shifted back, blurring lineage boundaries. Collectively, both SC and TC drive secretory cells toward squamous/keratinizing programs and weaken antimicrobial defenses. High-tar conventional smoke (TC) produces quantitatively stronger remodeling-particularly of squamous-metaplasia and mucin-repair programs-than low-tar slim smoke (SC), reinforcing the conclusion that TC imposes greater disruption on the compromised COPD secretory compartment.

## Conclusion

In the present study, we established a translationally relevant air–liquid interface (ALI) model using primary airway epithelial stem cells derived from patients with COPD and systematically evaluated the cellular and molecular responses to high-tar conventional cigarette smoke (TC), low-tar slim cigarette smoke (SC), and filtered air (NC). Physiological assays demonstrated that both smoke regimens elevated IL-1β expression, compromised ciliary structural integrity, and increased intracellular reactive oxygen species, with TC consistently producing more pronounced changes than SC. Single-cell RNA sequencing resolved six major epithelial populations (basal, basal-suprabasal provisional, secretory, secretory-like provisional, ciliated, and stressed cells) and enabled high-resolution, cell-type-specific analyses.

hdWGCNA of ciliated cells identified distinct co-expression modules: NC-associated modules were enriched for cilium organization, assembly, and polarity programs; SC-associated modules reflected moderate adaptive and defense responses; and TC-associated modules indicated more extensive reprogramming of ciliary functional networks. In the basal lineage, both smoke conditions triggered thousands of differentially expressed genes, activation of oxidative stress, ER stress/UPR, and squamous metaplasia pathways, together with developmental regression toward a less-committed, stem-like state along the differentiation axis; quantitative comparison showed that TC produced an approximately 1.7-fold stronger regression effect than SC. Secretory and secretory-like provisional cells exhibited compression of the differentiation continuum, downregulation of antimicrobial gene programs, and a coordinated shift toward keratinization and squamous-oriented transcriptional programs, again more evident under TC exposure.

These observations are consistent with, and extend, previous single-cell studies of smoking effects on human airway epithelium. In vivo scRNA-seq of tracheal epithelium from smokers versus never-smokers has shown that cigarette smoke perturbs every epithelial cell type through both shared pan-epithelial programs and hundreds of cell-type-specific response genes, including shifts in secretory and ciliated populations[16]. Complementary ALI-based scRNA-seq experiments using small airway epithelial cells from healthy donors and COPD patients demonstrated that cigarette smoke expands inflammatory and squamous (KRT6A) basal cell states, alters basal-to-secretory differentiation trajectories, and elicits largely similar directional responses in COPD- and healthy-derived cultures[17]. Earlier bulk and comparative toxicology studies of conventional versus reduced-tar or low-tar cigarettes likewise reported differential cytotoxicity and gene-expression changes, generally with stronger biological activity associated with higher-tar products[18]. Our work builds on these foundations by providing a direct side-by-side comparison of TC versus SC within a COPD patient-derived ALI system, integrating physiological readouts with modular co-expression analysis (hdWGCNA) and differentiation-axis quantification that together highlight the relative magnitude of influence exerted by the two smoke formulations[19–20].

Collectively, the data indicate that high-tar conventional cigarette smoke exerts greater influence than low-tar slim cigarette smoke across inflammatory, structural, and lineage-specific molecular dimensions in already compromised COPD airway epithelium. These findings may help inform discussions surrounding continued smoking and product switching among COPD patients.

This study has several limitations. The analyses are primarily transcriptomic; although immunofluorescence and ROS assays provided initial phenotypic support, broader protein-level and functional validation of the identified modules and pathway changes remains limited. The exposure regimen models repeated acute rather than long-term chronic smoke exposure and therefore may not fully capture the cumulative dynamics of continued smoking in patients. Sample size was constrained by the number of available COPD donors, and the absence of parallel healthy-donor ALI cultures precludes direct dissection of disease-specific versus shared responses. Future multi-omics, longer-term exposure, and functional studies will be required to further refine the cell-type-specific mechanisms underlying the differential influence of distinct cigarette formulations on COPD airway epithelium.

## Methods

### Patient recruitment and bronchoalveolar lavage fluid collection

Airway epithelial stem cells derived from patients with COPD were used for in vitro culture. Cells were obtained from Shanghai Tenth People’s Hospital (Shanghai, China). Patients were enrolled between January 2023 and December 2024 from the Department of Respiratory Medicine. Eligible participants were clinically stable patients with COPD who met the diagnostic criteria of the Guidelines for the Diagnosis and Management of Chronic Obstructive Pulmonary Disease (2021 Revision), defined as a post-bronchodilator forced expiratory volume in 1 second to forced vital capacity ratio (FEV /FVC) of <0.70. Clinical stability was defined as stable or mild respiratory symptoms (cough, sputum production, and dyspnea) with recovery to the pre-exacerbation baseline for at least 4 weeks. Patients with concomitant bronchial asthma, active pulmonary tuberculosis, lung malignancies, autoimmune diseases, or a history of acute exacerbation or respiratory tract infection within the preceding 4 weeks were excluded.

Bronchoalveolar lavage fluid (BALF) was collected according to a standardized protocol. A flexible bronchoscope was introduced via the oral or nasal route and advanced to the target pulmonary segment. For localized lesions, lavage was performed in the corresponding segment; for diffuse disease, the right middle lobe or the lingular segment of the left upper lobe was selected. Sterile saline prewarmed to 37 °C (total volume 100–250 mL) was instilled in 3–5 aliquots of 25–60 mL each and recovered by gentle negative-pressure aspiration. An overall recovery rate of >40% was required (>30% for lower-lobe lavage). Samples were considered acceptable when the proportions of red blood cells and epithelial cells were <10% and <5%, respectively. Ethical approval for this study was granted by the Institutional Review Board of Shanghai Tenth People’s Hospital (Approval No. 24K153), and all enrolled subjects provided signed informed consent.

### Culture and air–liquid interface differentiation of COPD patient-derived airway epithelial cells

Cells were expanded in PneumaCult™-Ex Plus Medium (STEMCELL Technologies, Cat. No. 05040). Upon reaching an appropriate density, cells were seeded onto 24-well Transwell inserts (Corning Costar, Cat. No. 3450; polyester membrane, 0.4 μm pore size). Air–liquid interface (ALI) culture was established using the PneumaCult™-ALI Medium Kit (STEMCELL Technologies, Cat. No. 05002). Under ALI conditions, the apical surface of the cells was exposed to air while the basolateral surface remained in contact with medium. Cells were maintained and differentiated at the ALI for 14–20 days until a stable, well-differentiated airway epithelial layer was formed, after which cigarette smoke exposure experiments were performed.

### Experimental groups and cigarette smoke exposure system

To compare the effects of different types of cigarette smoke on COPD patient-derived airway epithelial cells, three experimental groups were established: an air control group, a regular cigarette group, and a low-tar slim cigarette group, with three independent replicates per group. The regular cigarette group was exposed to LACX with a nominal tar yield of 11 mg/cigarette, whereas the low-tar slim cigarette group was exposed to LBXB with a nominal tar yield of 3 mg/cigarette. The air control group was exposed to air using the same exposure apparatus and for the same duration as the cigarette smoke groups. Cigarette smoke was generated using an RM20S smoking machine (Germany). For exposure, 24-well Transwell inserts containing differentiated airway epithelial cells were placed in the smoke exposure chamber, allowing the apical surface of the cells to be directly exposed to cigarette smoke or air. Throughout the exposure period, the cells were maintained under ALI conditions.

Fresh PneumaCult™-ALI Medium was continuously supplied to the basolateral compartment, while the medium was simultaneously drained through the outlet into a waste reservoir. The inflow and outflow were adjusted according to the liquid level to maintain a dynamic balance, ensuring that the medium remained in contact with the basolateral side of the Transwell membrane without flooding the apical surface. This configuration maintained a stable air–liquid interface throughout the exposure period while continuously supplying nutrients to the cells. The medium flowed in a single-pass manner and was not recirculated, thereby providing continuous basolateral liquid and nutrient support and mimicking a continuously perfused basolateral environment.

### Puffing and smoke dilution conditions of the RM20S smoking machine

The smoke method of the RM20S smoking machine was set to Full Puff, R&D. The puff volume was 35 mL, with a puff duration of 2.0 s and a piston sweep time of 1.9 s. The puff frequency was set to one puff per minute, corresponding to a puff period of 60 s and an intermission of 58.0 s. The correction factor for puff volume was set to 1.000. For smoke dilution, Stages 0, 1, and 2 of the RM20S dilution program were activated. The target volumes for Stages 0, 1, and 2 were set to 70, 50, and 100 mL, respectively. The remain volumes for Stages 1 and 2 were both set to 10 mL. The exhaust time was set to 5 s. The same puffing protocol and smoke dilution parameters were applied to both the regular cigarette and low-tar slim cigarette groups to ensure comparable exposure conditions.

### Repeated cigarette smoke exposure protocol

Each smoke exposure session used 12 cigarettes, with five puffs taken from each cigarette, resulting in a total of 60 puffs per exposure session. Cells in each experimental group underwent two rounds of exposure. Following the first exposure, the Transwell inserts were removed from the exposure chamber and transferred to an incubator at 37 °C with 7.5% CO for a 3-h recovery period. The cells were then subjected to a second exposure under the same smoking protocol, dilution parameters, and cigarette consumption as those used in the first exposure. The air control group underwent the same experimental procedures and incubation schedule but was exposed to air instead of cigarette smoke. After completion of the second exposure, cells were returned to the incubator and maintained at 37 °C with 7.5% CO for an additional 4 h to allow the cellular responses to cigarette smoke exposure to develop. The cells were then immediately subjected to subsequent physiological and phenotypic analyses.

### Assessment of cellular phenotypes and physiological responses after cigarette smoke exposure

Following two rounds of exposure and a subsequent 4-h recovery period, airway epithelial cells from the different treatment groups were assessed for oxidative stress, inflammatory responses, mucus production, and cilia-associated phenotypes. Intracellular reactive oxygen species (ROS) were detected by fluorescence staining using a ROS Assay Kit (Biosharp, Cat. No. BL714A) to evaluate cigarette smoke-induced oxidative stress. Cell nuclei were counterstained with DAPI.

After completion of the exposure protocol, the Transwell membranes containing airway epithelial cells were collected, fixed, paraffin-embedded, and sectioned for subsequent immunofluorescence staining. For IL-1β immunofluorescence staining, tissue sections embedded in paraffin underwent baking at 60 °C for 1 h, xylene deparaffinization, and rehydration through a descending ethanol gradient. Antigen unmasking was achieved by microwave-assisted heating in pH 9.0 EDTA buffer for 15 min. After cooling and three consecutive PBS washes (5 min each), slides were permeabilized with 0.3% Triton X-100 for 15 min and blocked with 5% normal goat serum for 30 min at room temperature. Primary antibody incubation against IL-1β (R&D Systems, catalog no. AF-201-SP; 1:100 dilution) was performed overnight at 4. After three washes with PBST for 10 min each, the sections were incubated with YSFluor™ 594 Donkey Anti-Goat IgG (H+L) secondary antibody (Cat. No. 34312ES60; 1:500) for 1 h at room temperature in the dark. Following three additional washes with PBST for 10 min each, the sections were mounted using an antifade mounting medium containing DAPI and imaged under a fluorescence microscope. MUC5AC and α-Tubulin double immunofluorescence staining was performed to evaluate mucus-related and cilia-associated phenotypes, respectively. MUC5AC was used as a marker of mucus-associated epithelial phenotypes, whereas α-Tubulin was used to visualize ciliary structures. The primary antibodies used were anti-MUC5AC (Abcam, Cat. No. ab3649; 1:100) and anti-α-Tubulin (Abcam, Cat. No. ab179484; 1:1000). All experimental groups were processed using identical staining procedures, image acquisition settings, and image analysis conditions.

### Single-cell RNA sequencing library preparation and sequencing

Cells recovered from ALI cultures of the NC, SC, and TC groups were processed for single-cell RNA sequencing using the DNBelab C4 platform according to the manufacturer’s protocol. Briefly, single-cell suspensions were prepared, encapsulated, and subjected to library construction. Libraries were sequenced on an appropriate high-throughput platform to generate raw reads for downstream analysis.

### Quality control, normalization, and cell clustering

Raw gene-barcode matrices were imported into R and analyzed with the Seurat package. Cells with outlier values for tnFeature_RNA, nCount_RNA, or percentage of mitochondrial transcripts were excluded. The remaining high-quality cells were normalized, followed by graph-based clustering and visualization by uniform manifold approximation and projection (UMAP). When necessary, integration procedures within Seurat were applied to mitigate batch effects across biological replicates [21]. Cell clusters were annotated based on the expression of canonical lineage markers (including KRT5 and TP63 for basal cells, SCGB1A1 and MUC5AC for secretory cells, FOXJ1, TPPP3 and DNAH5 for ciliated cells) together with additional functional markers.

### Differential expression analysis

Differentially expressed genes (DEGs) between exposure groups (SC versus NC, TC versus NC, and TC versus SC) within each annotated cell type were identified using the Wilcoxon rank-sum test implemented in Seurat. Genes with adjusted P-value < 0.05 and |log fold-change| ≥ 0.25 were considered significantly differentially expressed.

### Gene Ontology enrichment analysis

Gene Ontology (GO) biological-process enrichment of DEGs or module-associated genes was performed with the clusterProfiler package using the org.Hs.eg.db annotation database [22]. Significantly enriched terms were defined by conventional adjusted P-value and q-value thresholds and were simplified where appropriate to reduce redundancy.

### HdWGCNA analysis

hdWGCNA was applied specifically to the Ciliated cell population. Metacells were constructed to reduce data sparsity, co-expression networks were built, and modules were detected. Module eigengenes were calculated and compared across NC, SC, and TC conditions. Hub genes within each module were identified and visualized as interaction networks. Module genes were subsequently subjected to GO enrichment analysis as described above.

### Module scoring, developmental axis projection, and pseudobulk quantification

Functional module scores (including Club secretory identity, Squamous metaplasia, Inflammation NFκB, Oxidative stress NRF2, and ER stress/UPR) were computed with Seurat’s AddModuleScore function using curated gene sets. For developmental trajectory analysis, commitment axes for the basal and secretory lineages were defined from NC cells using the top differentially expressed genes between opposing poles of each lineage; all cells were then projected onto these axes to quantify shifts in differentiation state. Pseudobulk expression values (log counts per million) for selected marker genes were calculated per donor and condition to support quantitative comparisons across exposure groups.

---

**Figure S1.**
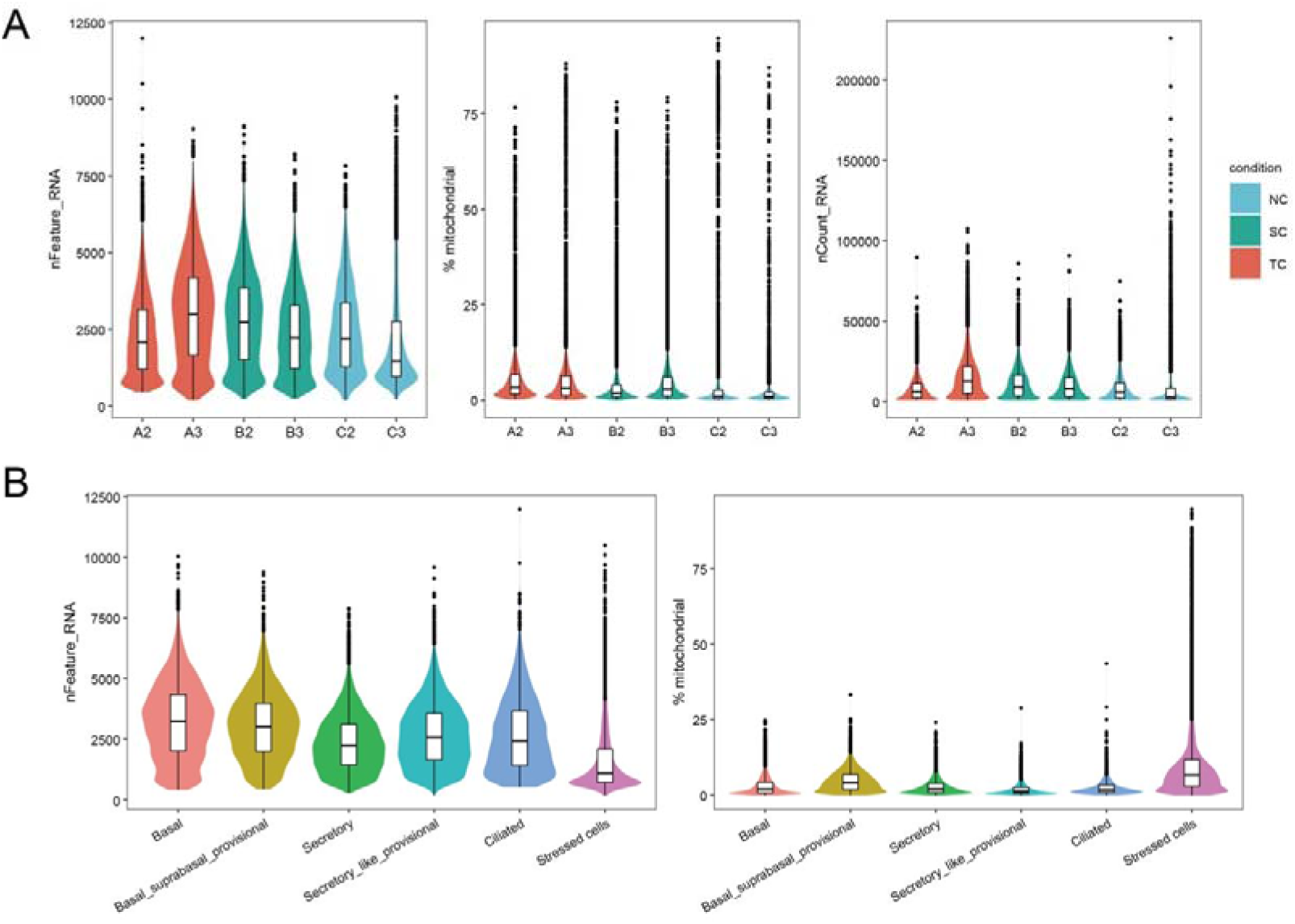
Quality control metrics and sequencing depth evaluations for single-cell RNA sequencing libraries. (A) Violin plots displaying the distribution of quality control parameters, including nFeature_RNA (left), mitochondrial gene percentage (% mitochondrial, middle), and total molecule counts (nCount_RNA, right), across individual experimental samples. (B) Violin plots showing the distribution of nFeature_RNA (left) and % mitochondrial (right) mapped across each annotated cell cluster, demonstrating uniform quality control thresholds across all cell populations.

## Declarations

### Author contributions

Jialei Jiang, Yushan Tian, and Yinuo Zhao performed the experiments, analyzed the single-cell transcriptomic data, and drafted the manuscript. Chen Tan, Xiao Li, and Hongjuan Wang contributed to sample collection, experimental validation, and substantive revision of the manuscript. Longjiang Fan and Hongyu Chen conceived, designed, and supervised the study. All authors read and approved the final manuscript.

### Data availability

The datasets generated during the present study will be publicly released upon final acceptance of the manuscript.

### Ethics approval

Not applicable.

